# Multifractality Nature of Microtubule Dynamic Instability Process

**DOI:** 10.1101/2020.05.28.121749

**Authors:** Vahid Rezania, Ferry C. Sudirga, Jack A. Tuszynski

## Abstract

The irregularity of growing and shortening patterns observed experimentally in microtubules reflects a dynamical system that fluctuates stochastically between assembly and disassembly phases. The observed time series of microtubule lengths have been extensively analyzed to shed light on structural and dynamical properties of microtubules. Here, for the first time, Multifractal Detrended Fluctuation analysis (MFDFA) has been employed to investigate the multifractal and topological properties of both experimental and simulated microtubule time series. We find that the time dependence of microtubule length possesses true multifractal characteristics and cannot be described by monofractal distributions. Based on the multifractal spectrum profile, a set of multifractal indices have been calculated that can be related to the level of dynamical activities of microtubules. We also show that the resulting multifractal spectra for the simulated data might not be comparable with experimental data.

**Statement of Significance:** Microtubules are some of the most important subcellular structures involved in a multitude of functions in all eukaryotic cells. In addition to their cylindrical geometry, their polymerization/depolymerization dynamics, termed dynamic instability, is unique among all protein polymers. In this paper we demonstrate that there is a very specific mathematical representation of microtubule growth and shrinkage time series in terms of multifractality. We further show that using this characteristic, one can distinguish real experimental data from synthetic time series generated from computer simulations.

## Introduction

In a dynamical system, complexity arises when its subunits interact non-linearly with each other. Existence of a critical threshold, irregularity and sudden phase transitions are most common features of many complex systems. Among many biological systems with intrinsic complexities, dynamic instability of microtubules (MTs) can be ranked very high as it has been the subject of numerous investigations for decades [1-3].

Microtubules are cylindrical-shaped polymers of tubulin (a family of globular proteins) found in the cytoplasm of eukaryotic cells and are a component of the cytoskeleton [4]. Microtubules are crucially involved in many cellular activities, including cell divisions (meiosis and mitosis), trafficking, and motility. Not surprisingly, dysfunctionalities of microtubules are correlated with a number of diseases, including various cancers, Alzheimer disease, and Parkinson disease [5]. Tubulin proteins that make up a microtubule assemble and disassemble themselves spontaneously in a stochastic manner, leading to a non-equilibrium process commonly referred to as microtubule’s dynamic instability [6].

Microtubules are formed by polymerization of its subunits, namely *α* and *β* tubulin heterodimers, into a cylindrical form with a hollow space inside which is called the lumen. In humans and most mammals, several distinct isotypes of *α* and β tubulins have been identified and their respective amino acid sequences mapped. Isotype variations within the *β* tubulin class are rather well understood in vertebrates where there are six main isotypes of *β* tubulins (I, II, III, IV, V, and VI) expressed by separate genes. These different *β* isotypes have been associated with different functional roles in eukaryotic cells [7]. Rezania et al [8] listed some of the key biophysical properties of various human *β*-tubulin such as their electrostatic charge, dipole moment, volume, surface area as well as specifically the properties of their C-terminal tails which are crucially important for both protein binding and microtubule assembly processes. Interestingly, even though these physical characteristics of *β* tubulin isotypes are quite similar, microtubule stochastic behaviour seems to change notably depending on its *β* tubulin isotype composition [8].

Dynamic instability of microtubules is driven by a competition between assembly and disassembly states of tubulin dimers at the microtubule ends. The so-called plus-end (distal) is more dynamic than the minus-end (proximal). Assembly, or growth, requires the presence of two GTP (Guanine Tri-Phosphate) nucleotides bound to an *αβ* heterodimer subunit. Tubulin-bound GTP is prone to hydrolyzation into GDP (Guanine Di-Phosphate) when bound to the *β* monomer while it does not hydrolyze when bound to the *α* -monomer. GDP-bound tubulin dimers are prone to disassemble. GTP-bound tubulins at the growing end of the microtubule appear to protect the entire structure from disassembly forming a so-called GTP-cap, as the microtubule’s inner GDP-bound tubulins do not seem to spontaneously depolymerize. When GTP hydrolysis catches up to the tip of the microtubule, however, the entire structure is prone to a rapid disassembly called a catastrophe [10].

Earlier statistical studies on microtubule data have mainly focused on analyzing length histograms of microtubules in order to shed light on their probability distributions [8,11]. As shown in these studies, histograms of MT length distributions have an exponential long tail and a peak corresponding to relatively short microtubules that suggests a characteristic of Poissonian processes. Furthermore, the resulting inverse proportionality of the frequency of catastrophes with respect to the growth velocity of a microtubule can be interpreted as an indication of the protective presence of a GTP cap at the growing end of a microtubule [12].

Recent studies employed different statistical methods for modeling stochastic signals such as compressed sensing, maximum likelihood, and hidden Markov models (HMMs) to capture the underlying signal variability [13]. Compressed sensing has been used to reconstruct signals using fewer samples than the Nyquist rate. The maximum likelihood technique has been applied to estimate parameters for a linear dynamic system from the observed data. Many wavelet-based hidden Markov processes have been also involved to capture real-world non-Gaussian signals [14].

Among the non-Gaussian variations, fractal behaviors have been found in biomedical signals from a wide range of physiological phenomena such as ECG, EEG, MR, and X-ray images [15]. Identifying scale-invariant variations not only can be used to differentiate between healthy and pathological conditions, but also to distinguish various types of unhealthy conditions. Biomedical time series such as interspike interval of neuron firing, stride interval of human walking, inter-breath interval of human respiration, and interbeat intervals of the human heart have all shown scale-invariant characteristics that can serve as a diagnostic tool [16-20]. In addition, scale invariant structures have also been found in spatial phenomena such as the branching of lungs, the vascular system and the nervous system [21-24], bone structure [25], and their properties such as the fractal dimension are able to differentiate between healthy and cancer tissues [26]. Furthermore, it has been proposed that changes in the scale-invariant structure of biomedical signals may result from the adaptability of physiological processes to new conditions [27]. As a result, a successful treatment of pathological conditions might change disturbed fractal structure and improve health. Therefore, fractal analysis of data structures can be used as a promising prognostic and diagnostic tool in biomedical signal processing.

Furthermore, a common theoretical approach to study stochastic properties of MTs is based on either the two-state model with growing-shrinking, or the three-state model with growing-pause-shrinking [1-3]. Effectively, such models are analogous to a 1D drift-diffusion random walk, where the drift velocity, *v*_drift_, and diffusion constant, *D*, depend on the four main microscopic parameters: growth (*v*_+_), shrinkage (*v*_−_), catastrophe (*f*_c_) and rescue (*f*_r_) rates:

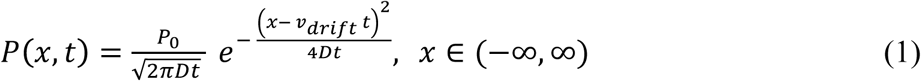

where *P*(*x,t*) is the probability density function and

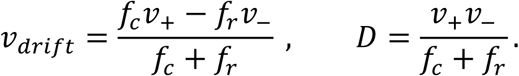

As a result, *<x>* = *v*_drift_ *t* and *<x*^2^*>* = *v*^2^_drift_ *t*^2^ + 2*Dt* ∼ *t*^2^ at large *t* [28]. Many physical and mathematical models have been proposed to shed light on the MT dynamic instability. Recently, Jonasson et al. [29] proposed a new model by introducing two different critical tubulin concentrations to provide a more unified explanation for many experimental observations of MT dynamic instability. Using their results, one can calculate the equivalent four parameters of the two-state model, i.e. (*v*_+_, *v*_−_, *f*_c_, *f*_r_). In addition, Cassimeris et al. [30] have also simulated an array of MTs using a Monte Carlo model to study the role of phase transitions and tubulin concentration on MT dynamic instability and MT length distribution. In a completely different approach, Rezania and Tuszynski studied MT polymerization and depolymerization processes by considering a semi-classical method that is often used in quantum mechanics [31]. They showed that the dynamics of a MT can be mathematically expressed via a 3D cubic-quintic nonlinear Schrodinger equation.

In this paper, however, we aim to investigate whether dynamic instability processes involving microtubules can be modelled using a fractal stochastic approach. We implement this approach on both experimental data, using *in-vitro* raw data primarily obtained from [8] as well as simulated data generated by methods described in [29], [30] and [31]. This provides a new comparison methodology between experiments and simulations.

Here, we adopted the *multifractal detrended fluctuation analysis* (MFDFA) method described in [32] to perform a statistical analysis of MT length data to shed light on the dynamics occurring during polymerization/depolymerisation processes. In addition, we used MT data assembled from two different isotypically-purified *β*-tubulins, namely *αβ*_II_ and *αβ*_III_ in order to understand how different isotype compositions may affect MT dynamics.

Fractal signals are usually divided into two categories: monofractal and multifractal structures. In general, the monofractal structures are those whose variations can be scaled by a single power law exponent that is independent of time and space. However, spatial and temporal variations often appear in the scale-invariant structure of a biomedical signal. As a result, these types of data cannot be scaled by a single exponent. These spatial and temporal variations indicate a multifractal structure of the biomedical signal that is defined by a multifractal spectrum of power law exponents. The multifractal spectrum identifies the deviations in fractal structure within time periods with large and small fluctuations [32-34]. Such variabilities have been observed in heart rate signals where the width and shape of the spectrum can also differentiate between patients with heart diseases such as ventricular tachycardia, ventricular fibrillation and congestive heart failure [35-36]. The multifractal structure analysis of EEG and series of interspike intervals has also been able to differentiate between the neural activities of brain areas [18].

Figure1 shows selected experimental time series data for the lengths of microtubule isotypes *αβ*_II_ (top panel) and *αβ*_III_ (middle panel). The lower panel demonstrates a simulated uniformly random data set over the same time sample as MT data for comparison purposes. By a simple observation from the following sample data one can conclude that:

**Figure 1.**
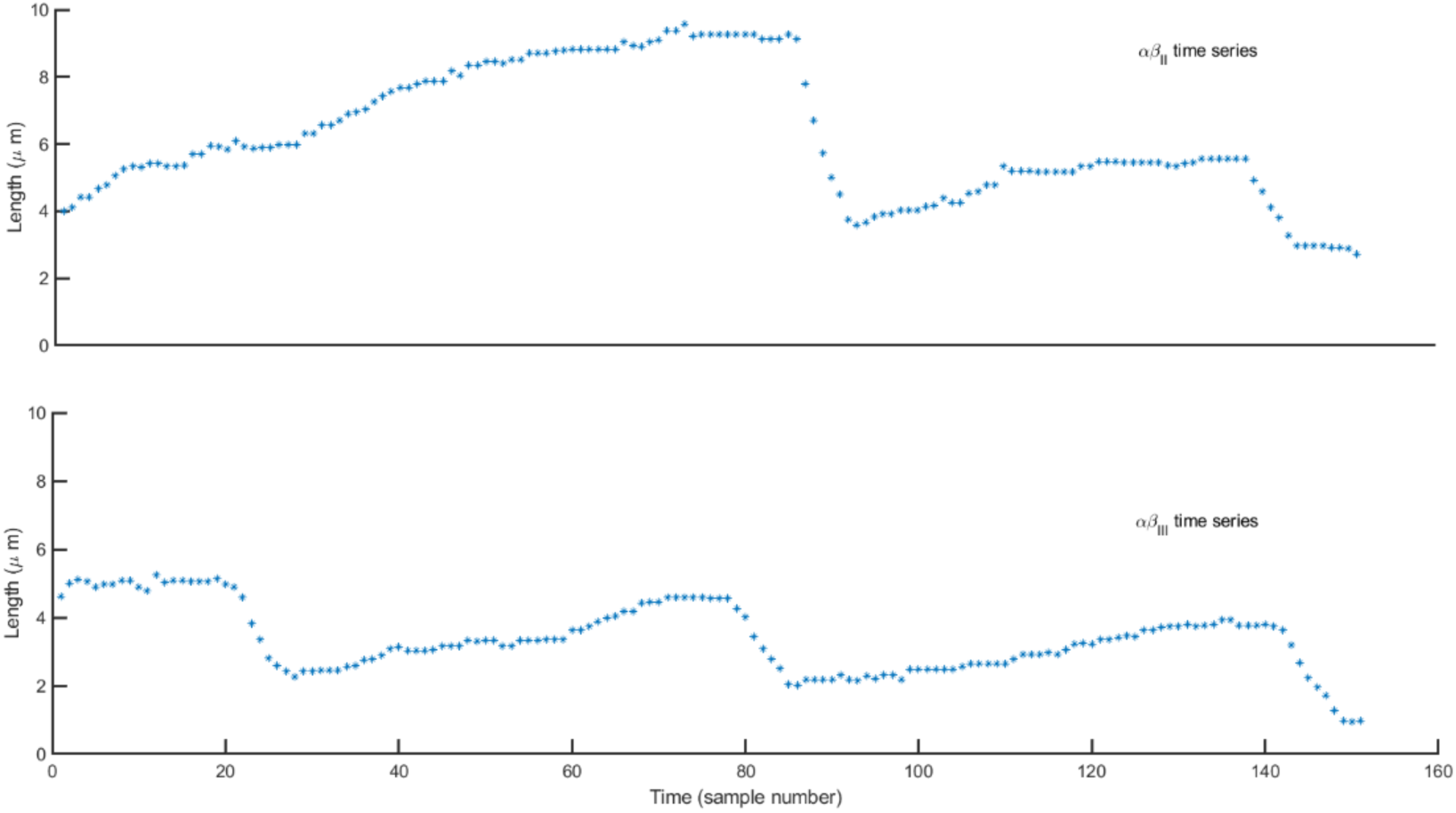
Selected sample data of the growth pattern of microtubule isotypes *αβ*_II_ (top panel) and *αβ*_III_ (lower panel) from [8].

i. microtubules usually spend most of their time in the growing phase rather than in the shortening phase [6,8].
ii. changes in length are trending, suggesting the possibility of a positive serial correlation [6].
iii. the shortening phase can be fast and drastic.
iv. volatility of length changes does not seem to be constant; i.e. volatility clustering and relatively frequent sharp jumps are observed.

Hence, modelling the stochastic behaviour of microtubules requires a different framework and might not be captured fully by the 1D drift-diffusion random walk models. One can explain the aforementioned behaviour by using a model that involves fractality, a pattern of self-affinity that shows up both at larger and smaller scales. Some fractals are deterministic, such as the Mandelbrot set and the Koch snowflake that are defined by iterative equations [37]. In Euclidian geometry, the length of an object can be defined as the limit of a sequence of approximations. For example, if we construct a polygon and increase the number of its sides, the length of the polygon converges to a limit, which is the circumference of the circle surrounding it. However, in fractal geometry, this might not be the case. Consider the Koch snowflake, which is made of an infinite number of self-affine structures. The perimeter length of the snowflake converges to an infinite number. Empirically, fractality can be quantified by the fractal dimension *d*_F_, which is a measure of fractal “roughness of shape” [37].

Fractals can also be stochastic. For example, almost no two snowflakes are the same. One measure of stochasticity in fractals is to calculate the Hurst exponent *H* introduced by Hurst [38], see the Methods section. For noisy time series the Hurst exponent will be in the interval between 0 and 1 whereas for a random walk type time series, it will be above 1. A Hurst exponent of 0.5 indicates a purely random process, or white noise. That means that a corresponding time series has an independent or short-range dependent structure. A process with *H* > 0.5, however, implies a positive persistence or a correlation between the past trend and future behavior. This means that if a variable was increasing in the past, it should continue to increase. In such a process, time series differences have some tendency to trend together: positive movements are more likely to be followed by positive movements and *vice versa. H* < 0.5 indicates a negative persistence or anti-correlated behavior resulting in a dampening process. The closer the *H* value is to 0, the rougher or more jagged the trend looks like.

Hence, a fractal model with *H* > 0.5 can explain the trending patterns of microtubule growth and why microtubules tend to spend more time in the growing phase [6]. One relatively simple example of a fractal model is the Fractional Brownian Motion developed by Mandelbrot and van Ness [39].

Fractal analysis can capture some behaviors observed in microtubule data but not all. Specifically, the volatility clustering behavior observed in the microtubule data can only be explained if the stochastic process is multifractal. This is a more complex multi-layered fractal structure (i.e. fractals of fractals), in which different segments of the time series exhibit localized behaviors, driven by different *H* values for different statistical moments [40]. This is unlike the Fractional Brownian Motion process, in which *H* is constant for all statistical moments, i.e. a monofractal stochastic process. A model data set from a multifractal stochastic process is charted below. Note that the cumulative time series exhibits localized patterns that are similar to the longer series and that volatility ebbs and flows throughout the time series.

## Methodology

By its very nature, empirical data that come from a (hypothesized) multifractal process would be non-stationary and many conventional statistical procedures cannot be applied to such data. Kantelhardt et al. [41] pioneered a rather ‘computationally friendly’ method called the Multifractal Detrended Fluctuation Analysis (MFDFA) that has been used extensively by subsequent researchers [32, 42]. This research utilizes MFDFA algorithms as configured by [32]. The MFDFA method for a time series {*x*_*i*_}, i= 1, …, *N* proceeds as follows:

1. Integrate the time series to create the profile:

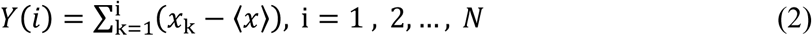

where ⟨*x*⟩is the average value of {*x*_*i*_}.
2. Divide the profile *Y*(*i*) into *N*_s_=*N/s*, non-overlapping segments of equal length *s*. Then, calculate the local trend for each segment by fitting the data using a least-squares fit, y_v_ (*i*), with either linear, quadratic, or higher polynomial orders.
3. Detrend each segment *v* = 1, 2, …, *N*_s_ by subtracting the best fit, y_*v*_ (*i*), from the data. Then compute the mean-squared variance:

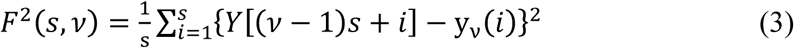

where y_*v*_ (*i*) is the polynomial fitted to the segment *v*(*i*).
4. Average all segments to obtain the *q*-order fluctuation function

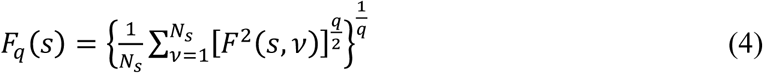

where *q* is a real number. It is interesting to note that for positive *q, F*_*q*_(*s*) measures large fluctuations, while for negative *q, F*_*q*_(*s*) measures small fluctuations in data.
5. Determine the localized generalized Hurst exponent *H*_*q*_ at each *q*-order by repeating the above calculation to obtain the fluctuation function *F*_*q*_(*s*) for a different box size *s*:

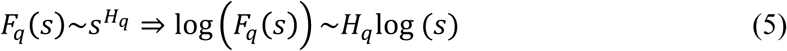

Hence, the generalized Hurst exponent, *H*_*q*_, is the fluctuation parameter that describes the correlation structures of the time series at different magnitudes and can be empirically quantified by running a regression of log *F*_*q*_(s) vs log *s* for each *q*-order. In practice, quantification becomes more inaccurate as *q* becomes more positive or negative. Here, *H*_*q*_ is calculated for -5 ≤ *q* ≤ 5, as suggested in [32]. Note that *H*_2_ (i.e. the Hurst exponent at *q* = 2) is identical to the basic Hurst exponent of a monofractal stochastic process [6, 43].

Note that the value of the Hurst exponent at the zeroth order, *H*_0_, cannot be determined by using Eq. (4) because of the diverging exponent. The logarithm averaging procedure should be used [41, 44]:

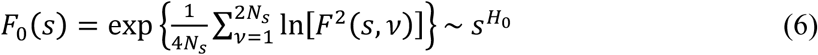

Information contained in the relationship between *H*_*q*_ and *q* can be further transformed into useful statistical parameters as defined below [41]:

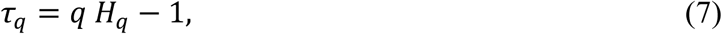

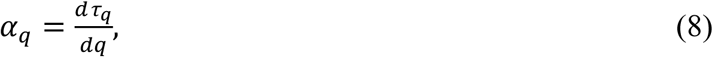

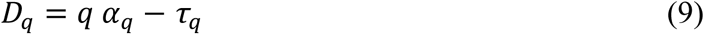

Here, *τ*_*q*_ is called the *q*-order mass exponent and can be used to quantify the scaling properties of the data. *α*_*q*_ is the *q*-order singularity exponent that represents local Hurst exponents and *D*_*q*_ is the *q*-order dimension or the singularity spectrum. Each local Hurst exponent, *α*_*q*_, quantifies the local singular behavior and thus relates it to the local scaling of the series. The plot of *D*_*q*_ as a function of *α*_*q*_ is also called the multifractal spectrum. For white noise time series, the mass exponent *τ*_*q*_ in Eq. (7) is a linear function of the *q*-order. This linear relationship of *τ*_*q*_ leads to a constant *α*_*q*_, the first derivative of *τ*_*q*_, as expected (see Figure 3). The nonlinear dependence of the mass exponent *τ*_*q*_ with the *q*-order, however, signifies the multifractality in the time series that can be decomposed into many subsets exhibiting different local Hurst exponents, *α*_*q*_. The resulting multifractal spectrum will be usually a large arc where the difference between the maximum and minimum *α*_*q*_ is called the multifractal spectrum width, also called *the degree of multifractality, Δα*_*q*_ = max(*α*_*q*_) – min(*α*_*q*_), see Figure 3. The multifractal spectrum width is an important parameter; a wider spectrum implies more intense volatility clustering and extreme jumps in data values, something that Mandelbrot [45] referred to as wilder randomness. Conversely, in the random walk systems, the multifractal spectrum width is much smaller than in the multifractal systems, congruent with their properties of constant volatility [32].

**Figure 2.**
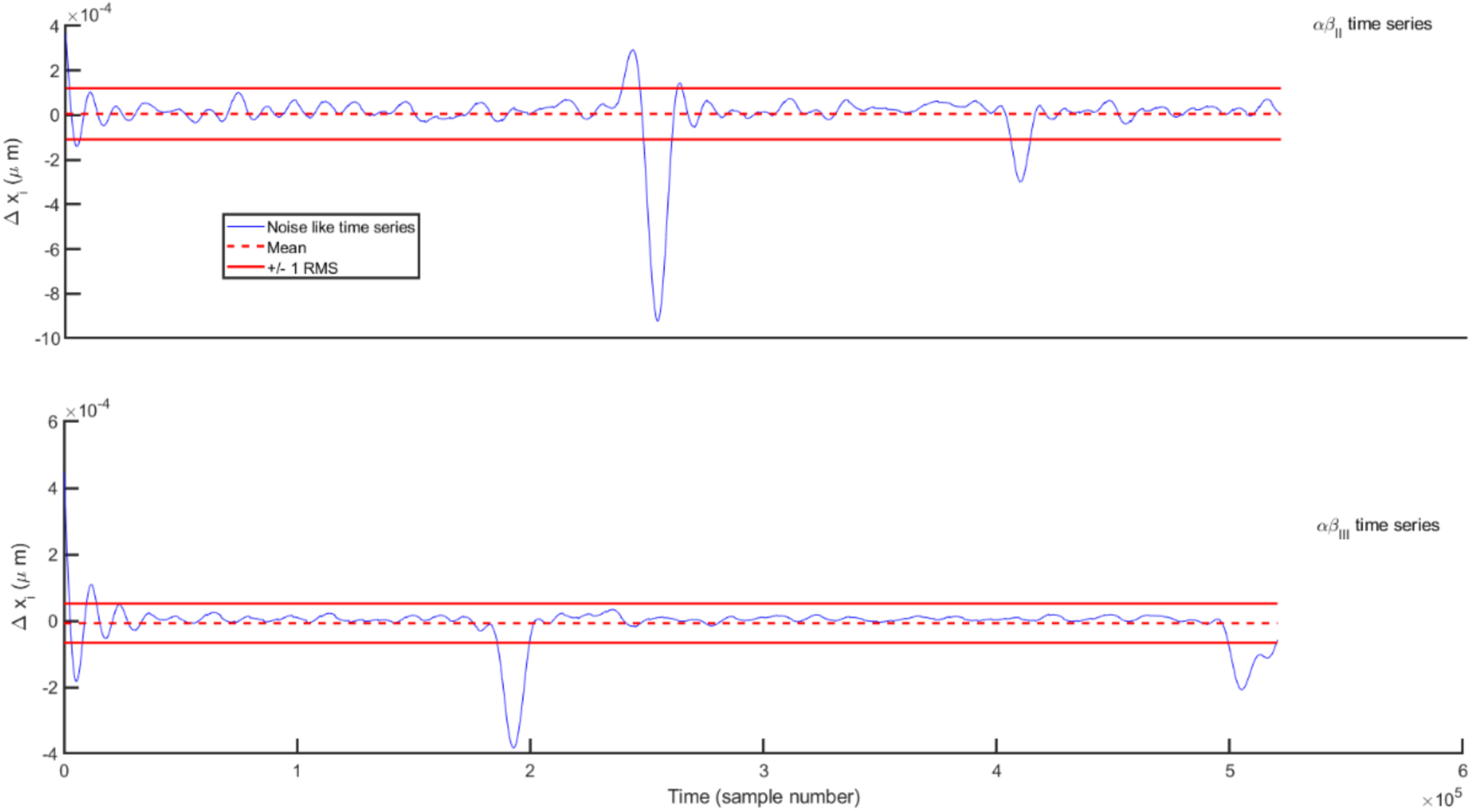
The difference time series, Δ*x*_i_ = *x*_i+1_ − *x*_i_, *i* = *2*, …, *N. αβ*_II_ (top panel), *αβ*_III_ (middle panel), and white noise (lower panel) with zero average (red dashed lines) and ±RMS (red solid lines). The data are plotted in terms of the sample number rather than time for better comparison.

**Figure 3.**
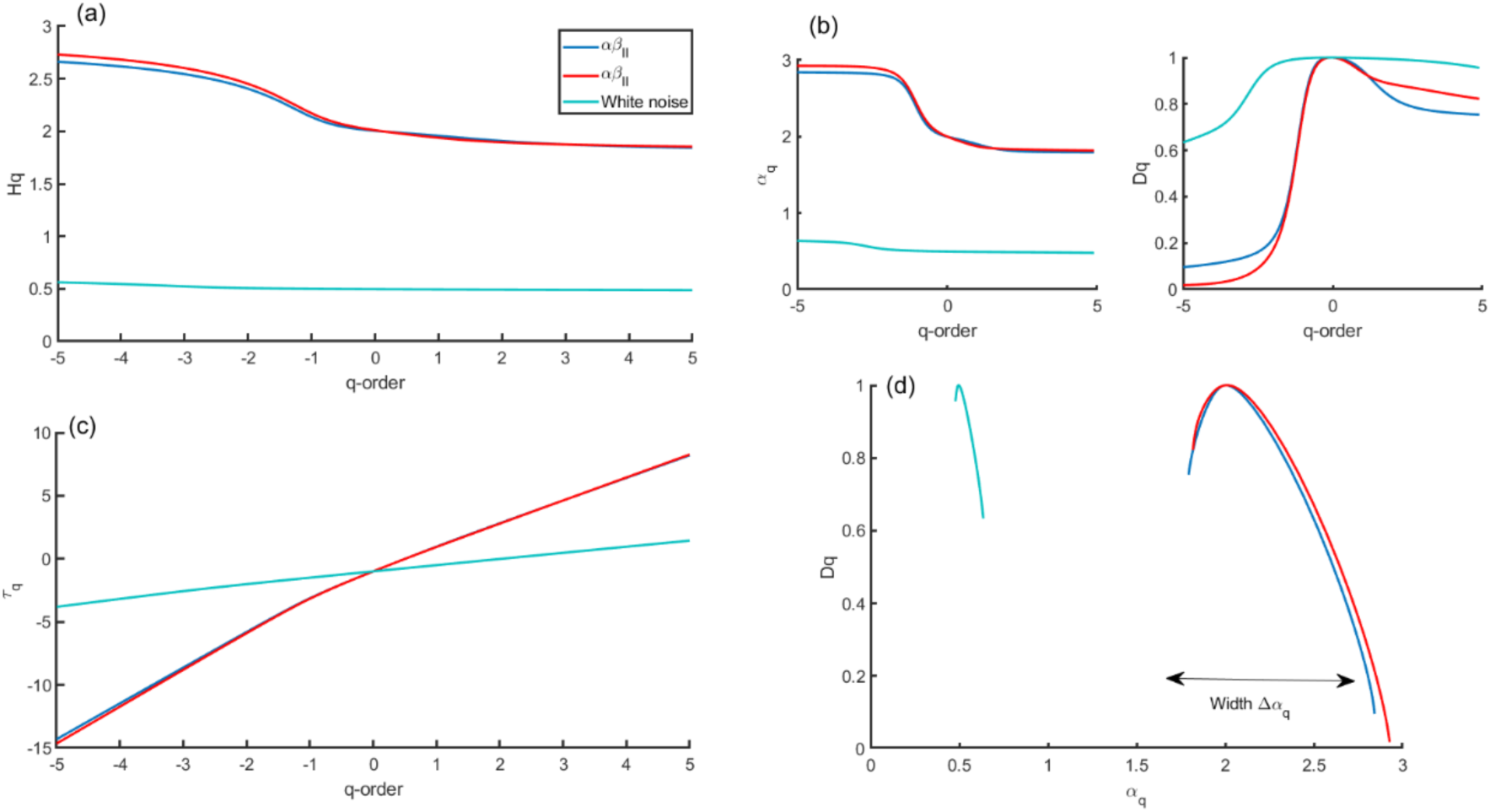
Multiple representations of the multifractal spectrum for the selected *αβ*_II_, *αβ*_III_ and the simulated white noise time series. (a) *q*-order Hurst exponent *H*_*q*_ (b) mass exponent *τ*_*q*_ (c) *q*-order multifractal spectrum of *D*_*q*_ and *α*_*q*_ (d) multifractal spectrum of *D*_*q*_ as a function of *α*_*q*_. The arrow indicates the difference between the maximum and minimum *α*_*q*_, which is called the multifractal spectrum width.

In general, the multifractality in time series arises either from: (i) fat-tailed probability distributions of increments, compared with a Gaussian distribution, or from (ii) different long-range temporal correlations for small and large fluctuations [42, 46]. To identify the contribution of either of these sources, two procedures of shuffling and phase randomization have been used in the literature. These procedures are usually employed to study the degree of complexity of time series to distinguish different sources of multifractality in the time series. The type (i) can be tested by phase randomized surrogates that weaken the non-Gaussian effects in the data. The procedure preserves the amplitudes of the Fourier transform but randomizes the Fourier phases that eliminate nonlinearities while keeping only the linear properties of the original time series [46]. In the shuffling procedure, the values are put in random order. As a result, the type (ii) can be tested by shuffling as it erases the long-range temporal correlation for small and large fluctuations and conserves the overall distribution of the data. In other words, through the shuffling procedure the distribution function of the original data remains exactly the same but without memory. This leads to the *q*-order singularity spectrum, *D*_*q*_, reaching a maximum at *α*_*q*_ ∼ 0.5. Furthermore, if multifractality in the time series is due to a broad distribution function, *α*_*q*_ obtained by the surrogate method will be independent of *q*. However, if both kinds of multifractality are present in time series, the shuffled and surrogate series will show a weaker multifractality strength than the original one [41]. Here, we apply these methods to MT time series data and compare with the original data.

## Results and Discussions

### Experimental data

Here, we studied experimental microtubule data based on tubulin samples extracted from bovine brains as described in [8]. There are 26 data sets of *αβ*_II_ and 30 data sets of *αβ*_III_ tubulin isotypes. It is necessary to note that the availability of samples was limited, which constrains our analysis. In general, the growing and shortening dynamics data for individual MTs at their plus ends were recorded by differential interference contrast video microscopy. Data points representing microtubule lengths were collected as a time series with 2-6 s intervals for as long as 10 minutes at a time. This means each microtubule data set only provided 120-250 data points. As suggested by Ihlen [32], however, ideally more than 1,000 points per sampling might be needed for robustness of the analysis.

Due to limited sample sizes, we increase the number of sampling rates by using Matlab resampling utility that implements an anti-aliasing lowpass filter. It should also be noted that sampling rates were uneven (i.e. time intervals between data points are not fixed), possibly due to some manual snap-shooting of microtubule lengths during the experiments.

As stated previously, Figure 1 demonstrates a selected raw experimental length data of microtubule isotypes *αβ*_II_ (top panel) and *αβ*_III_ (middle panel). For comparison purposes, uniformly random (Gaussian) data over the same time sample as microtubule data have been generated (lower panel). Resampled data have not been shown.

Figure 2 shows the difference data, Δ*x*_*i*_ = *x*_*i*+1_ − *x*_*i*_, *i* = *2*, …, *N*, for each data set in Figure 1 after the resampling procedure. The results are plotted in terms of the sample number rather than time for better comparison. The root-mean-squared (RMS) value is only sensitive to differences in the amplitude of variation and not differences in the structure of variation. Notice the extreme amplitudes in both *αβ*_II_ and *αβ*_III_ results that are 3-4 times of their RMS values in comparison with the white noise with no such changes. As expected, these extreme changes are due to catastrophes in the corresponding data sets.

To utilize the multifractality of the data set, the multifractal spectrum is plotted and describes the singularity *D*_*q*_ as a function of *α*_*q*_. The typical procedure to graph the multifractal spectrum in the MFDFA method is to first convert multifractal Hurst exponent *H*_*q*_ to the *q*-order mass exponent *τ*_*q*_, and then convert *τ*_*q*_ to the *q*-order singularity exponent (or local Hurst exponent) *α*_*q*_, and finally to *q*-order singularity dimension *D*_*q*_ [41], see Figure 3. To determine the multifractal Hurst exponents for different orders *q*, the minimum window length of 5 data points that varies in the range of 10–20 s was used. The maximum window length that we used here is *int*(*N*/5) where *N* denotes the total number of data points in each data set. A total of 20 successive windows have been chosen that sets the length of each successive window increases by a factor of ∼ 2^1/8^ [32]. The multifractal Hurst exponents, *H*_*q*_, were determined by varying *q* in the range, −5 to +5 for all three data sets.

Figure 3 shows the *q*-dependence of *H*_*q*_, *α*_*q*_, *τ*_*q*_, and *D*_*q*_ for the selected *αβ*_II_ and *αβ*_III_ data sets as well as the simulated white noise for comparison purposes. As expected, the white noise time series depicts a linear relationship of the mass exponent *τ*_*q*_ with *q*-orders. The nonlinearity shape of *τ*_*q*_ reveals a multifractal behavior of the time series. For a monofractality feature that appears, the function *τ*_*q*_ would be a linear function of *q*-orders with a constant slope. The linear *q*-dependence of *τ*_*q*_ leads to a constant *α*_*q*_ that reduces the multifractal spectrum to a small arc for the white noise time series (see Figure 3d). Similar behavior to that for white noise data is also expected for a monofractal data set. Furthermore, the average value of the Hurst exponent for the white noise is about 0.52, which is expected and implies a *short-range* dependence in the time series. The mass exponents *τ*_*q*_ for the *αβ*_II_ and *αβ*_III_ data sets, however, show a curved *q*-dependence and, consequently, a decreasing singularity exponent *α*_*q*_. The resulting multifractal spectra represent a large arc where the difference between the maximum and minimum *α*_*q*_ is called the multifractal spectrum width, Δ*α*_*q*_, see arrow in Figure 3(d). Figure 4 shows the generalized Hurst exponent and multifractal spectrum for all *αβ*_II_ and *αβ*_III_ data sets. The Hurst exponent for both *αβ*_II_ and *αβ*_III_ data sets converges to ∼ 1.5, which signifies the non-Gaussian behavior of the data sets. In general, the Hurst exponent indicates for a time series by how fast the overall RMS of local fluctuations RMS grows with an increasing segment sample size (i.e., scale). The fast changing fluctuations in the time series will influence the overall RMS, for segments with small sample sizes (i.e., small scale) whereas slow changing fluctuations will influence it for segments with large sample sizes (i.e., large scale). The larger Hurst exponent, *H*, is usually translated as more slowly-evolving variations or a more persistent structure in multifractal time series. Furthermore, the Hurst exponent greater than 0.5 describes a *long-range* dependence and memory effects on all time scales according to the level of persistence. As *H* approaches 1 or higher, the time series becomes increasingly periodic [42].

**Figure 4.**
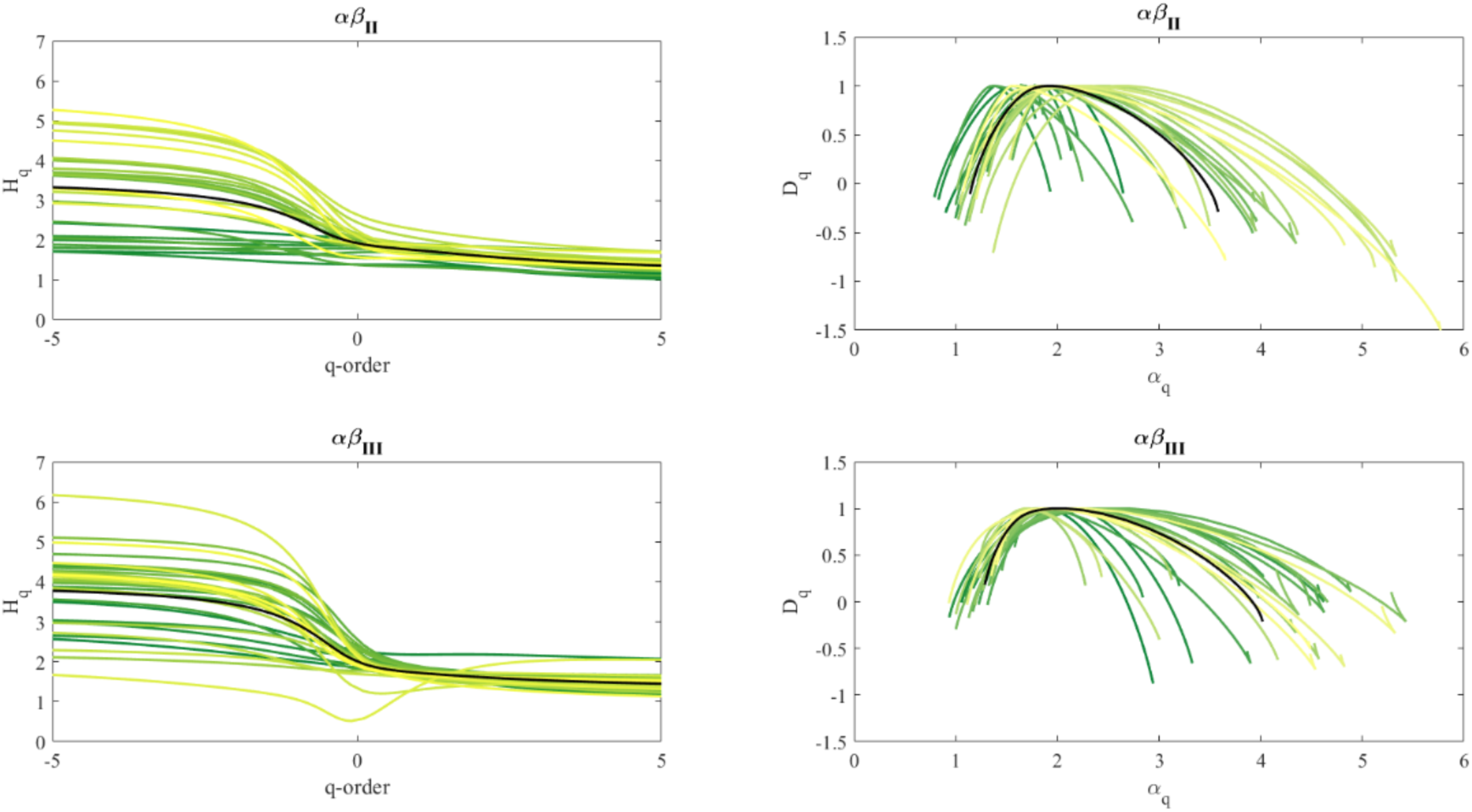
*q*-order Hurst exponent *H*_*q*_ (left panels) and multifractal spectrum of *D*_*q*_ as a function of *α*_*q*_ (right panels) for all available *αβ*_II_ and *αβ*_III_ data sets. The black line represents the average over corresponding data.

Figures 3 and 4 show that even though both *αβ*_II_ and *αβ*_III_ data sets show multifractality characteristics, there are still subtle differences between them. To measure the differences among these spectra, a set of multifractality indices has been calculated. These indices can also be related to the level of dynamical activities of microtubules. Following [42], the first index derived for a multifractal spectrum is the *degree of multifractality* of the spectrum, Δ*α*_*q*_

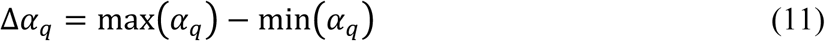

The wider *Δα*_q_ corresponds to the richer and stronger multifractality in data. On average, the degree of multifractality for *αβ*_II_ and *αβ*_III_ spectra is found to be 2.441 and 2.731, respectively.

Furthermore, in an ideal case, the shape of the multifractal spectrum, *D*_*q*_, is symmetric. However, for real-world data, this is not necessarily symmetric. The asymmetry in a spectrum is related to the volatility of the data set and is likely to be a genuine result and not merely a data or methodology artifact [47]. The left side of the multifractal spectrum *D*_*q*_ is affected by the positive *q* values, which filter out larger fluctuations, and the vice versa. As a result, asymmetry in *D*_*q*_ reveals non-uniformity, indicating that the corresponding large and small-scale fluctuations are driven by different mechanisms [47]. If the spectrum is symmetric, it means there is a balance between the mechanisms that determine each side of the spectrum. The equilibrium will be broken when one of the fluctuations (small or large) is dominant.

The *degree of asymmetry, A*, or the skewness in the shape of the spectrum can be calculated as [42]:

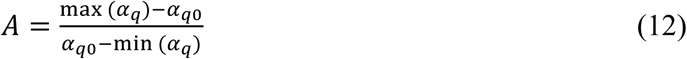

where *α*_*q*0_ is the value of *α*_*q*_ when *D*_*q*_ reaches its maximum. This spectrum is symmetric if *A* = 1, left-skewed for 0 < *A* < 1, and right-skewed when *A* > 1. For the *αβ*_II_ and *αβ*_III_ spectra, the skewness values are found to be 2.114 and 2.749, respectively. On average, the *αβ*_III_ isotype series appear to have a bolder right-skewed asymmetry (relatively more segments of low volatility than segments of high volatility) while the *αβ*_II_ isotype series appear to have a modest left-skewed asymmetry, see Figure 4.

In addition, the multifractal spectrum can also have either a left or right truncation [32]. The truncations basically measure a leveling of the *q*-order Hurst exponent for negative or positive *q* values, respectively. The truncation is estimated by the *singularity ratio C* that is defined by the ratio between 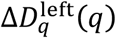 of the left side of the spectrum and 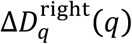 of the right side of the spectrum calculated in relation to the maximum fractal dimension, 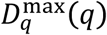:

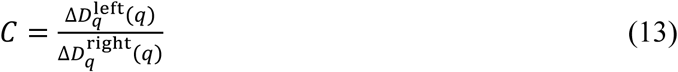

where

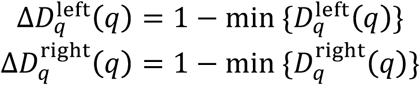

Here, min 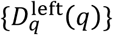 and min 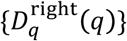 are minimum values of *D*_*q*_ in the left and right part of the spectrum, respectively. In this sense, for the singularity index *C* < 1, the right-hand side is truncated, while for *C* > 1 the truncation occurs on the left-hand side. Interestingly, a long left tail indicates that the time series has a multifractal structure that is insensitive to local fluctuations with small magnitudes. Conversely, a long right tail implies that the time series has a multifractal structure that is insensitive to local fluctuations with large magnitudes. For the *αβ*_II_ and *αβ*_III_ spectra presented here, the average singularity ratios are 0.8595 and 0.6736, respectively.

In order to identify the source of mutltifractality whether it is due to type (i) the broad (fat-tail) probability density function or type (ii) the long memory correlations in the small and large fluctuations, we calculated the multifractal indices for the original MT data sets and the resampled ones, the shuffled and surrogated data sets. Figure 5 demonstrates the averaged generalized *q*-order Hurst exponent and multifractal spectrum for the original (black curve), shuffled (red curve) and surrogated (blue curve) of *αβ*_II_ and *αβ*_III_ data sets. As shown, the multifractal spectrum *D*_*q*_ of *αβ*_II_ is fairly symmetric in comparison with that of *αβ*_III_. This suggests the large and small-scale fluctuations contribute more or less equally in *αβ*_II_ data. Furthermore, the result illustrates that both shuffling and surrogate applications reduce the multifractality strength of the original time series that demonstrates that both the long-range correlation and the fat-tail distribution contribute to multifractality. The multifractality is, however, much less pronounced in the shuffled time series than original and surrogated time series. As a result, one can say that the long-range correlations among fluctuations have major contribution in the observed multifractality of the MTs’ time series. This can also be seen by calculating various multifractality parameters that are summarized in Table 1. As shown, except for the degree of asymmetry, *A*, the (*h*_q0_, *H*_2_, *Δα*_q_, *C*)-quadruplets in the original time series are greater than their shuffled and surrogated partners, i.e. (*h*_q0_, *H*_2_, *Δα*_q_, *C*)_original_ > (*h*_q0_, *H*_2_, *Δα*_q_, *C*)_surrogated_ > (*h*_q0_, *H*_2_, *Δα*_q_, *C*)_shuffled_. The nonzero spectrum width of the shuffled time series, *Δα*_q_^shuffled^ > 0, shows a weak multifractal effect in the shuffled time series. This implies that while we have removed the correlations in the original series, the multifractality still remains in the shuffled time series. This highlights that the multifractality of MT data arises from both types with the main contribution from type (i), as the shuffled time series shows much less a pronounced multifractality than the original and surrogated time series [48-49].

**Table 1.** Multifractal indices for the original, shuffled and surrogated data.

| | $\alpha_{q0}$ | $H_2$ | Degree of multifractality, $\Delta\alpha_q$ | Degree of asymmetry, $A$ | Singularity ratio, $C$ |
| --- | --- | --- | --- | --- | --- |
| $\alpha\beta_{II}$ – original | 1.925 | 1.604 | 2.441 | 2.114 | 0.860 |
| $\alpha\beta_{III}$ – original | 2.016 | 1.617 | 2.731 | 2.749 | 0.674 |
| $\alpha\beta_{II}$ – shuffled | 0.605 | 0.534 | 1.083 | 3.152 | 0.297 |
| $\alpha\beta_{III}$ – shuffled | 0.560 | 0.476 | 0.990 | 3.763 | 0.270 |
| $\alpha\beta_{II}$ – surrogated | 1.567 | 1.376 | 1.132 | 1.630 | 0.654 |
| $\alpha\beta_{III}$ – surrogated | 1.503 | 1.307 | 1.645 | 2.623 | 0.526 |

**Figure 5.**
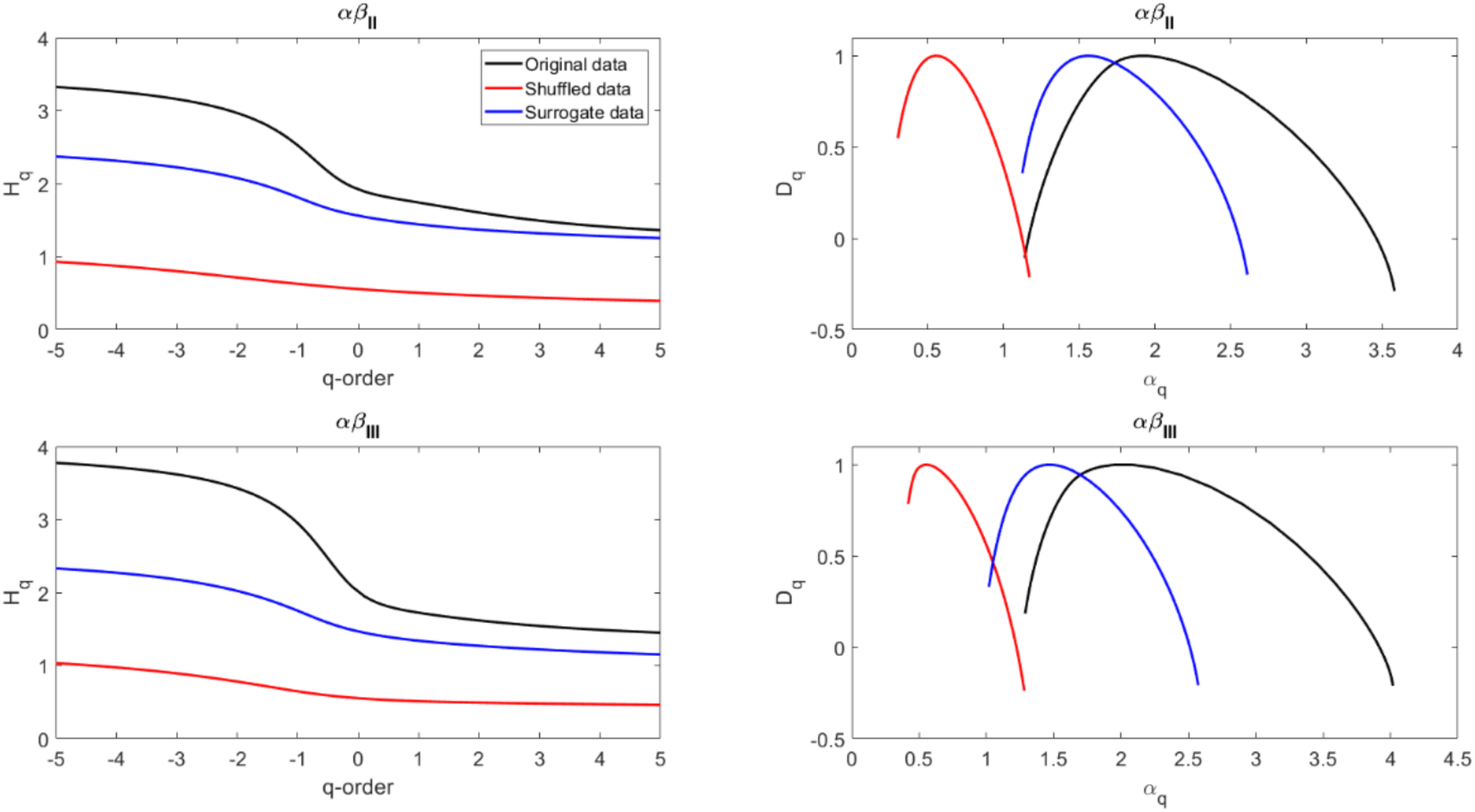
Averaged *q*-order Hurst exponent *H*_*q*_ (left panels) and multifractal spectrum of *D*_*q*_ as a function of *α*_*q*_ (right panels) for original, shuffled, and surrogated *αβ*_II_ and *αβ*_III_ data sets.

### Simulated data

The MFDFA algorithm has been applied to three differently simulated data for comparison purpose. The first simulated data is based on the model proposed by Jonasson et al. [29] (hereafter model A) who studied MT polymerization/depolymerisation processes by considering two different critical tubulin concentrations to provide a better explanation for experimental observations of MT dynamic instability. The second set of data is generated based on Cassimeris et al. [30] (hereafter model M) who simulated an array of MTs using a Monte Carlo model and studied the effect of phase transitions and free tubulin concentrations on MT dynamics instability and MTs’ length distribution. The computational codes were provided as a supplementary material. The third data set was generated based on the work of Rezania and Tuszynski [31] (hereafter model C) who studied MT polymerization and depolymerization processes by applying a semi-classical method that is often used in quantum mechanics. They showed that the dynamics of a MT can be mathematically expressed via a 3D cubic-quintic nonlinear Schrodinger equation.

In Figure 6, we plotted two time series of the normalized MT length over the sample number as an example for each models. The upper panels are constructed based on the model A, the middle panels are simulated based on model B and the lower panels are generated using the model C. For each data set, similar parameters are chosen as described in corresponding papers.

**Figure 6.**
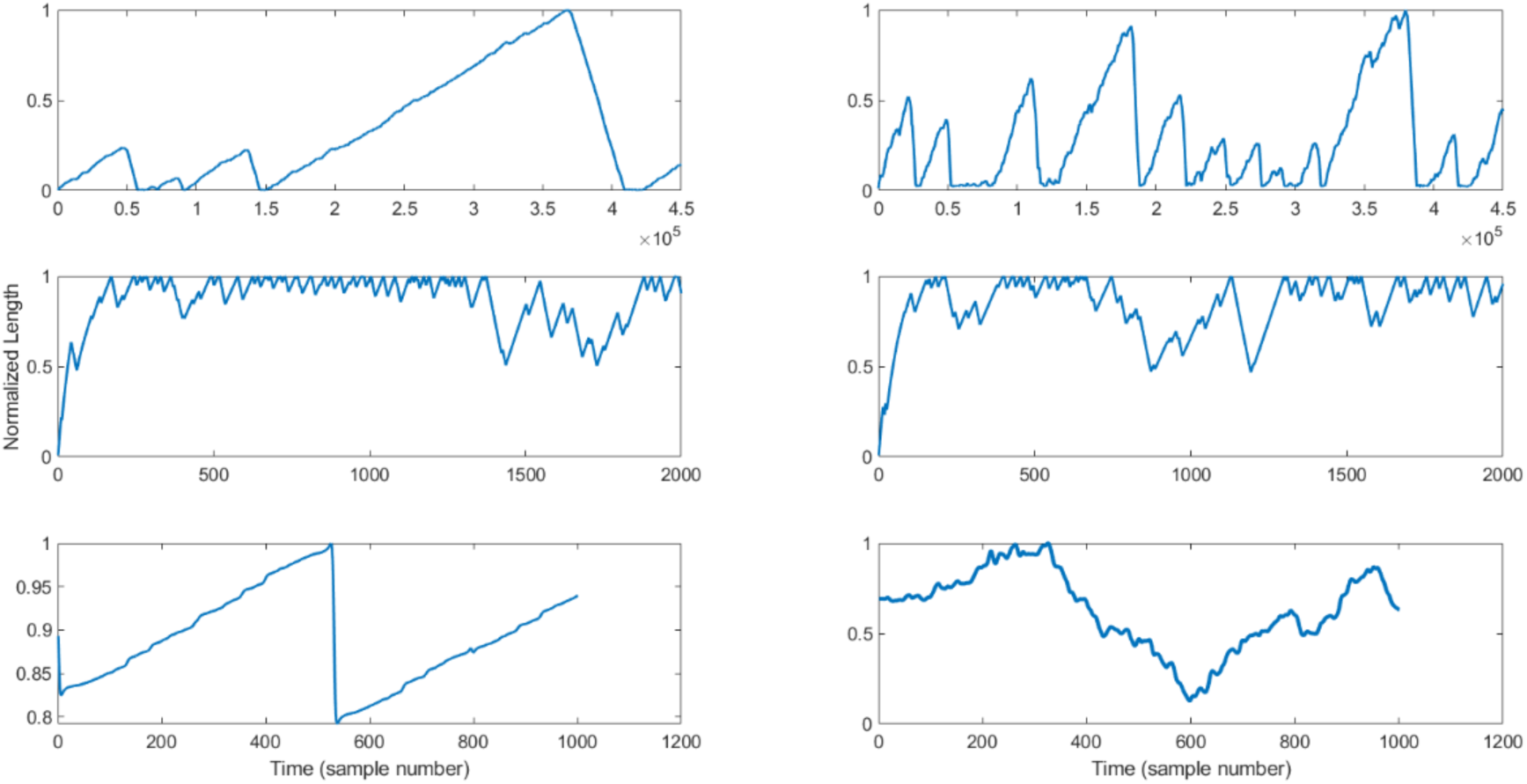
Simulated microtubule length data sets using three different methods described in models A (top panel), B (middle panel), and C (lower panel).

Similarly to experimental MT data sets, we performed the shuffling and the surrogated procedures on the simulated data from all three different models. Figure 7 shows the averaged generalized *q*-order Hurst exponent and multifractal spectrum for the original (black curve), shuffled (red curve) and surrogated (blue curve) of each simulated data set. Again, the upper, middle and lower panels are based on models A, B and C, respectively. Comparing Figures 5 and 7, the *q*-order Hurst exponent and multifractal spectrum, *H*_*q*_ and *D*_*q*_, for the simulated data in model C is more or less comparable with the experimental time series. Although the simulated time series from models A and B show some degrees of multifractality, their multifractal spectrums are sharply truncated from the right side of the spectrum. This means the large-scale fluctuations have very little contributions in their data constructions. This is obviously not the case for the simulated data in the model C.

**Figure 7.**
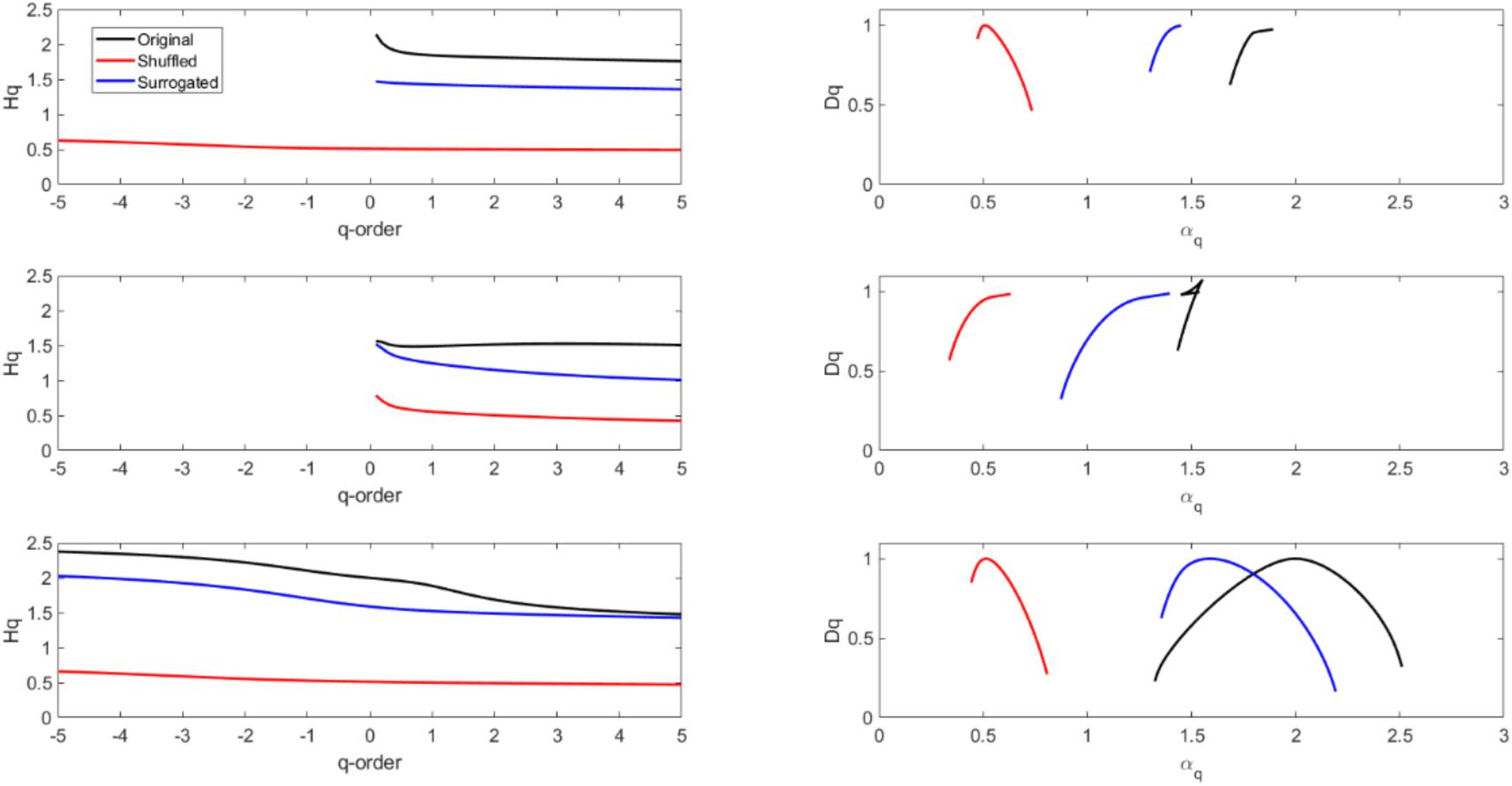
Averaged *q*-order Hurst exponent *H*_*q*_ (left panels) and multifractal spectrum of *D*_*q*_ as a function of *α*_*q*_ (right panels) for original, shuffled, and surrogated simulated microtubule data sets. Top panel: model A, middle panel: model B, and lower panel: model C.

## Conclusions

In this paper, we have used the MFDFA method to investigate the apparently stochastic nature of the experimental microtubule length data recorded in the course of 5–10 min intervals. These time series data describe the dynamic instability of microtubules that is driven by a competition between assembly and disassembly processes. In particular, we examined the data obtained for two different MT isotypes, namely *αβ*_II_ and *αβ*_III_, to compare their similarities and differences. Our results indicate that the microtubule growth data sets exhibit a multifractal characteristic: a nonlinear curve of *H*_*q*_ as a function of *q-*orders, as opposed to a constant *H*_*q*_ = 0.5 for all *q-*orders, which is characteristic of a random walk process, or a linear function of *q-*orders for a monofractal process. Furthermore, by comparing the generalized Hurst exponents of the original time series with the shuffled and surrogated ones, we have found that multifractality due to the broadness of the probability density function has a greater contribution than the long-range correlations. This may be interpreted as indicating that the MT polymerization dynamics is governed by different physical mechanisms at different time scales. For example, MTs start polymerizing from nucleation, which is a highly nonlinear process. This is then followed by a largely linear assembly process, which is interspersed with stochastic catastrophes and non-uniformly distributed rescues [50]. Therefore, on different time scales, different specific mechanisms dominate the MT dynamic instability process.

In addition, we have examined three different simulated data sets based on the models A, B, and C proposed by Jonasson et al. [29], Cassimeris et al. [30], and Rezania and Tuszynski [31], respectively. Models A and B generally lead to the known two-state/three-state model for MT dynamic instability with some additional information. Even though the time series fluctuations in all three models considered here can represent MT dynamic instability, see Figure 6, the multifractal analysis shows that their resulting spectrums are quite different. Both models A and B have shown a sharply truncated spectrum from the right side suggesting that the structure of the data is mainly due to small-scale fluctuations. Model C, however, has shown a more symmetric spectrum, similar to experimental data, confirming that both small- and large-scale fluctuations contribute more or less equally. See Figure 7.

Available microtubule data used in this research suggest the possibility that the microtubule dynamic instability process is stochastically multifractal. Due to the fact that our analysis was limited to two tubulin isotypes and the data sets were not sufficiently large, further research should be performed with longer time-series data beyond 10 minutes of microtubule observations in order to achieve greater statistical robustness. The evolution patterns of the multifractality and the geometrical properties of the visibility graphs would have further clinical usefulness to investigate patterns of polymerization time series for drug-treated microtubules and to derive causality relationships between microtubule stochastic behaviours with mechanistic effects at a molecular level.

## Acknowledgements

V.R. wishes to thank sincerely Dr. Goodson and her group as they provided the computational codes for this purpose and Philip Winter for his help in running some of the codes. V. R. also is grateful to MacEwan University for research grant for supporting this research. J.A.T. acknowledges with gratitude research support from NSERC (Canada) and the Allard Foundation.

